# Adaptation-induced oscillatory phototaxis and emergence of ordered density bands in the microswimmer *Chlamydomonas reinhardtii*

**DOI:** 10.1101/2024.06.09.598154

**Authors:** Zhao Wang, Alan C. H. Tsang

## Abstract

Biological microswimmers exhibit versatile taxis behaviors and switch between multiple behavioral states to navigate the environment and search for physiologically favorable regions. Here, we report a striking oscillatory phototaxis observed in *Chlamydomonas reinhardtii*, where cells swim back-and-forth under a constant, unidirectional light stimulus due to alternation between positive and negative phototaxis. This oscillatory phototaxis at the individual cellular level further leads to the emergence of a highly ordered, propagating band structure formed by high density *Chlamydomonas* cells collectively. We experimentally verify a unified phototaxis mechanism that couples light detection, light adaptation, flagella dynamics and cell reorientation, showing that transition between phototaxis modes is achieved by switching of flagella waveforms and modulation of flagella phase difference. Oscillatory phototaxis emerges as a semi-stable state in an overlapping light intensity regime for positive and negative phototaxis, where adaptation shifts the light intensity thresholds over times. This adaptation mechanism over multiple time scales enables phototactic microswimmers to effectively expand the survival range of light intensity and provide collective photoprotection for the colonies through the formation of dynamic band structures with high density.

---

Oscillatory motilities or switching between multiple behavioral states are common responses of biological cells to environmental stimuli: Neutrophils exhibit oscillatory motions in opposing gradients of chemoattractants to avoid being trapped in local chemical maxima [1, 2]; Copepods change swimming direction frequently with powerful jumps to escape from predators [3]; *Escherichia coli* exhibit run-and-tumble to search for food [4]. Such oscillatory motilities are uncommon for pond microbes’ responses to light (e.g., *Chlamydomonas reinhardtii, Euglena gracilis, Volvox carteri* and other green algae), which typically select between unidirectional positive and negative photo-taxis to swim towards and away from light, respectively [5–9]. This selection between different photoresponses enables these microswimmers to navigate favorable light environments for processes such as photosynthesis and population growth [10, 11], having profound implications to the sustainability of aquatic ecosystems.

## Oscillatory phototaxis and adaptation-induced density wave

Here, we report a striking oscillatory phototaxis observed in *Chlamydomonas reinhardtii* at the individual level, where the cells swim in oscillatory trajectories under a constant light stimulus from one side of the fluid chamber (Fig. 1a-c; Supplementary Movie S1, S2), in contrast to the unidirectional positive or negative phototaxis normally observed. After ⪆ 10 cycles which took *∼* 10 *−* 30 mins, the cells eventually transition to positive phototaxis and accumulate near the light source. We performed experiments in large fluid chambers (*∼* 4.5 cm *×* 2 cm in length and width, *∼* 300 *µ*m in depth) and verified that the oscillatory phototaxis was not due to boundary interactions or light reflection (Supplementary Section 2.2). This oscillatory phototaxis emerges under intermediate light intensities at 4000 *−* 8000 lux, lying between positive phototaxis under weak light (*∼* 150 lux) and negative phototaxis under strong light (*>* 15000 lux). While positive and negative phototaxis have been relatively well studied [12–20], there were only a few reports of similar oscillatory behaviors [21, 22]. More surprisingly, we discover that this oscillatory photo-taxis for individual cells leads to the emergence of ordered band structures formed by high density of *Chlamydomonas* cells collectively at the population level (Fig. 1d, Supplementary Movie S3). The density wave slowly propagates towards the light source at *∼* 100 *µ*m/min and reaches the end of the chamber after most cells transition to positive phototaxis due to adaptation. Yet, it is unclear how and why the oscillatory phototaxis emerges as a transition behavior, and what is the role of adaptation in this oscillatory behavior.

**Figure 1:**
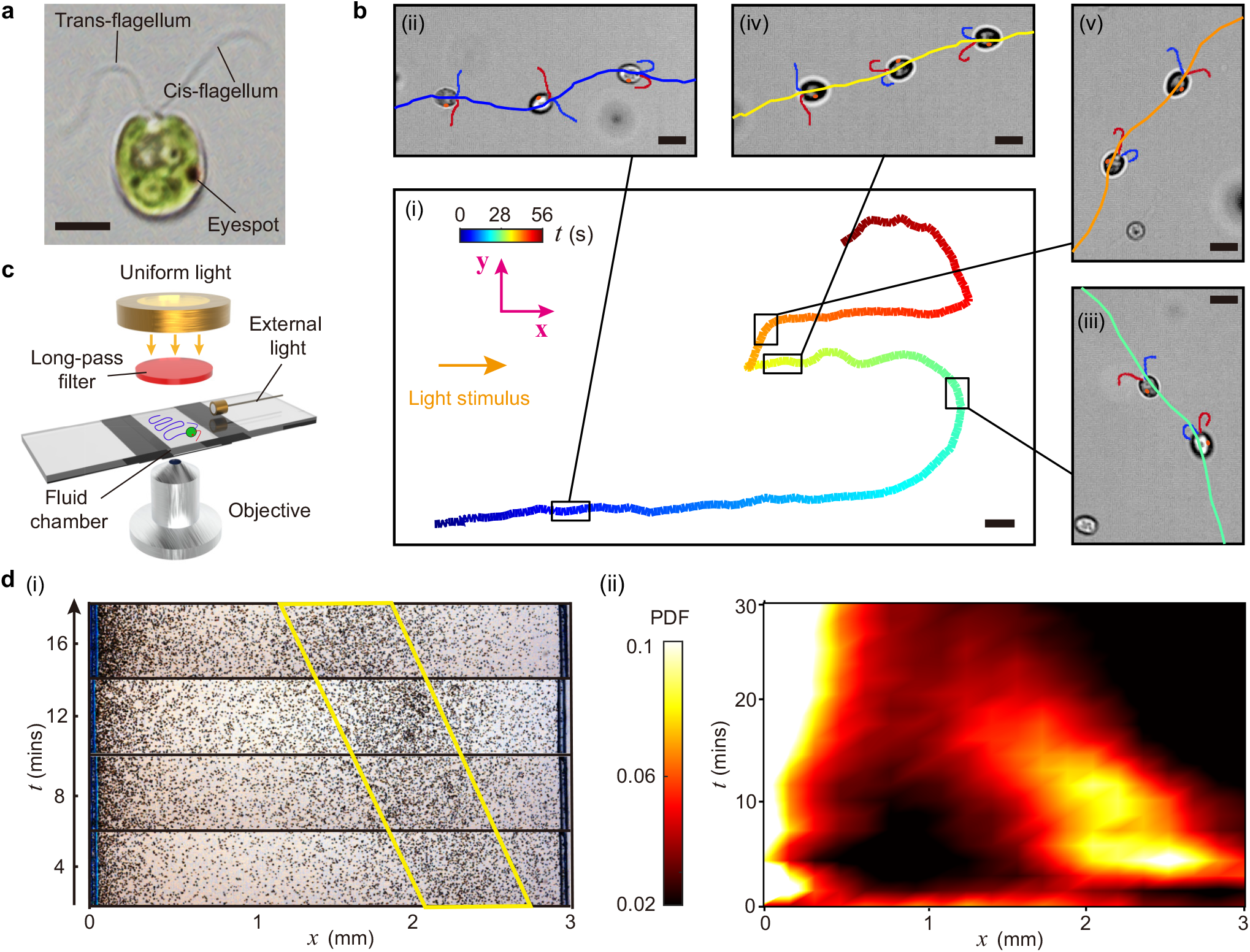
*Chlamydomonas reinhardtii* exhibit striking oscillatory phototaxis under a constant light field at intermediate intensity. **a**, *Chlamydomonas* has a red ‘eyespot’ (photosensory organelle) that converts light signals into phototactic responses of its cis- and trans-flagellum, located close to and far away from the eyespot, respectively. **b**, *Chlamydomonas* alternates between positive and negative phototaxis to exhibit oscillatory phototaxis. (i) A typical trajectory of oscillatory phototaxis. (ii-v) Overlaid time-lapse images of a cell at different phototaxis modes: (ii) equilibrium state of negative phototaxis, (iii) turning state of positive phototaxis, (iv) equilibrium state of positive phototaxis, (v) turning state of negative phototaxis. Cis- and trans-flagellum outlines are tracked and colored in red and blue, respectively. Eyespot is marked as an orange spot. Light stimulus aligns with 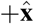 direction. **c**, A schematic of experimental set-up: cells in a fluid chamber are illuminated by red observing light from the top using a long-pass filter, which cells cannot detect [50]. An external white LED is applied from the side of the chamber. **d**, Emergence of density wave due to individual oscillatory phototaxis in a high density of *Chlamydomonas* cells. (i) Time-lapse images showing the propagation of density wave (highlighted by the yellow box). (ii) Kymograph of probability density function (PDF) of cells obtained by image thresholding. Scale bars: 5*µ*m (**a**), 200*µ*m (**b**(i)) and 10*µ*m (**b**(ii-iv)).

The phototactic microswimmer *Chlamydomonas reinhardtii* is a representative model organism for fundamental biological research including biosensing, photosynthesis, flagella dynamics and cell adhesion [23–26]. It also enables environmental and technological scientific discoveries such as bioremediation, bioconvection, food and biofuel production, optogenetics and cargo delivery [27– 31]. *Chlamydomonas* has a spheroidal shape with diameter of *∼* 10 *µ*m (Fig. 1a), swimming in (left-handed) helical trajectories at speed of *∼* 70 *−* 100 *µ*m/s while rolling around their long axis anticlockwise (observed from back) at 1-2 Hz [32]. *Chlamydomonas* has a red ‘eyespot’ that senses and converts light signals into coordination of *∼* 50-60 Hz beating patterns of its cis- and trans-flagellum, which enables the cell to steer its helical path in response to light and exhibit phototaxis. Although photoresponsive behaviors of *Chlamydomonas* have been extensively studied, the findings are mostly limited to characterization of cell motilities [13, 14, 33], flagellar dynamics for immobilized cells [15–18] and measurement of photoreceptor current [19, 20]. A systematic study connecting photoreception, adaptation, flagellar dynamics and phototaxis transition for free-swimming cells is lacking.

The novelty of the observed oscillatory phototaxis and the adaptation-induced density wave motivate our further investigation on their underlying biophysical mechanisms. In the following, we describe the basis of oscillatory phototaxis and its relationship to positive and negative phototaxis at the levels of individual cellular dynamics (Fig. 2) and subcellular flagellar beats (Fig. 3). We then introduce a photomechanical model that captures the observed oscillatory phototaxis (Fig. 4) and the adaptation-induced density wave (Fig. 5).

**Figure 2:**
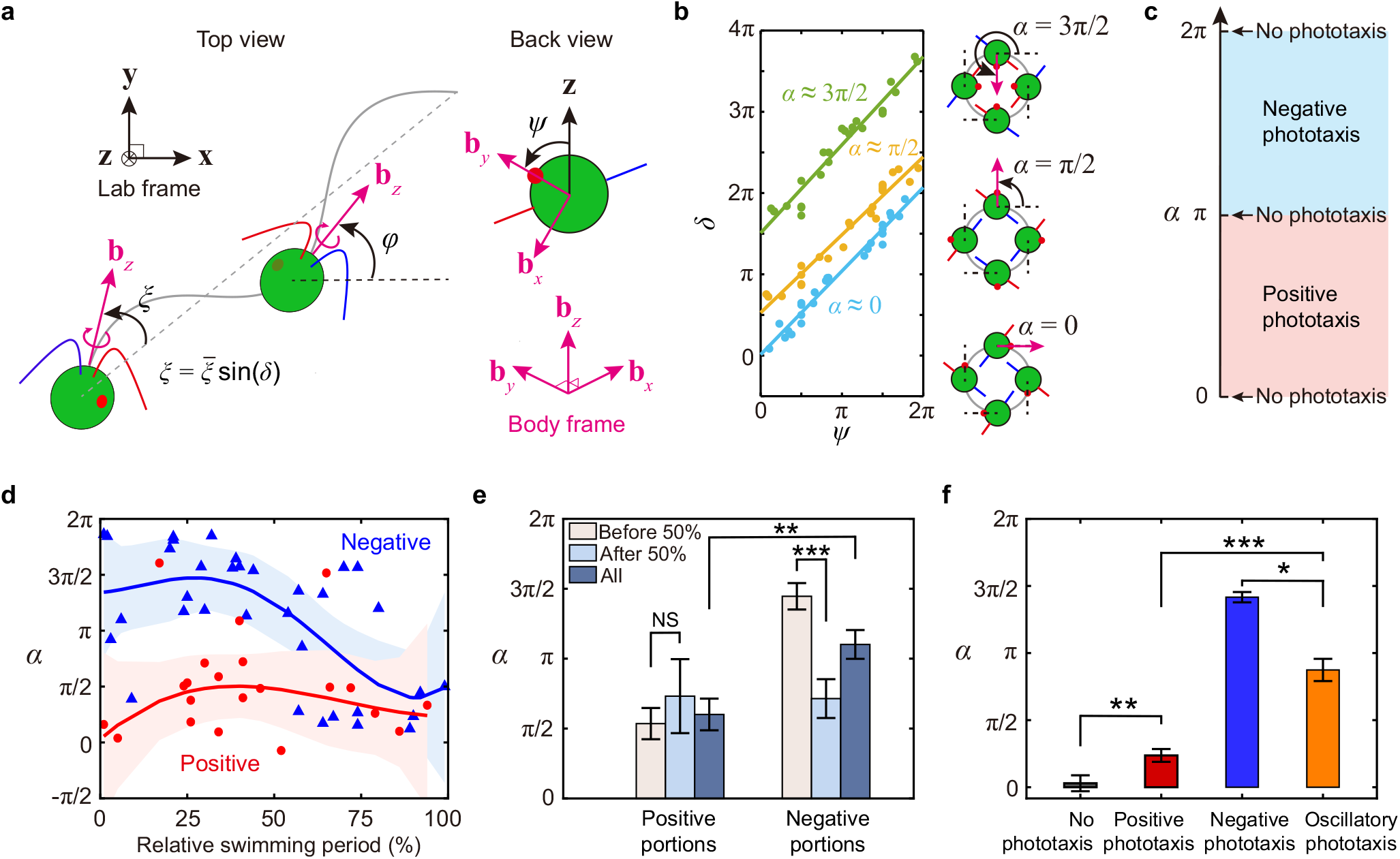
*Chlamydomonas* cells dynamically tune a phase angle *α* to switch between positive and negative phototaxis to enable oscillatory phototaxis. **a**, A schematic of parameter definitions for tracking the dynamics of *Chlamydomonas*. The cell swims in a helical trajectory and rotates along the body axis. Cis- and trans-flagellum are labeled in red and blue, respectively. **b**, Linear fit of the phase of the helical trajectory *d* and the roll angle *ψ* display a fixed phase relationship for cells under darkness (cyan, *α* = 0.04 *±* 0.13), positive phototaxis (yellow, *α* = 1.66 *±* 0.18) and negative phototaxis (green, *α* = 4.74 *±* 0.20). The phase angle *α* is defined as the intercept of the linear fit obtained by tracking cells for 3 roll cycles. **c**, A geometric model reveals that the phototaxis sign of *Chlamydomonas* is determined by the range of *α* (Supplementary Section 3). **d**, Changes in *α* during oscillatory phototaxis. The solid lines are obtained by spline interpolations, where the colored areas donate the 95% confidence interval. *n* = 34 samples in negative portions (blue color) and *n* = 21 samples in positive portions (red color) collected from *N* = 6 cells. **e**, Comparison of *α* between positive and negative portions before and after 50% of the relative swimming period as well as the whole swimming period. NS denotes no significant difference. **f**, Comparison of *α* between cells under darkness, positive phototaxis, negative phototaxis and oscillatory phototaxis, where *n* = 6 cells in each case. All error bars represent SEM throughout the paper if not stated otherwise.

**Figure 3:**
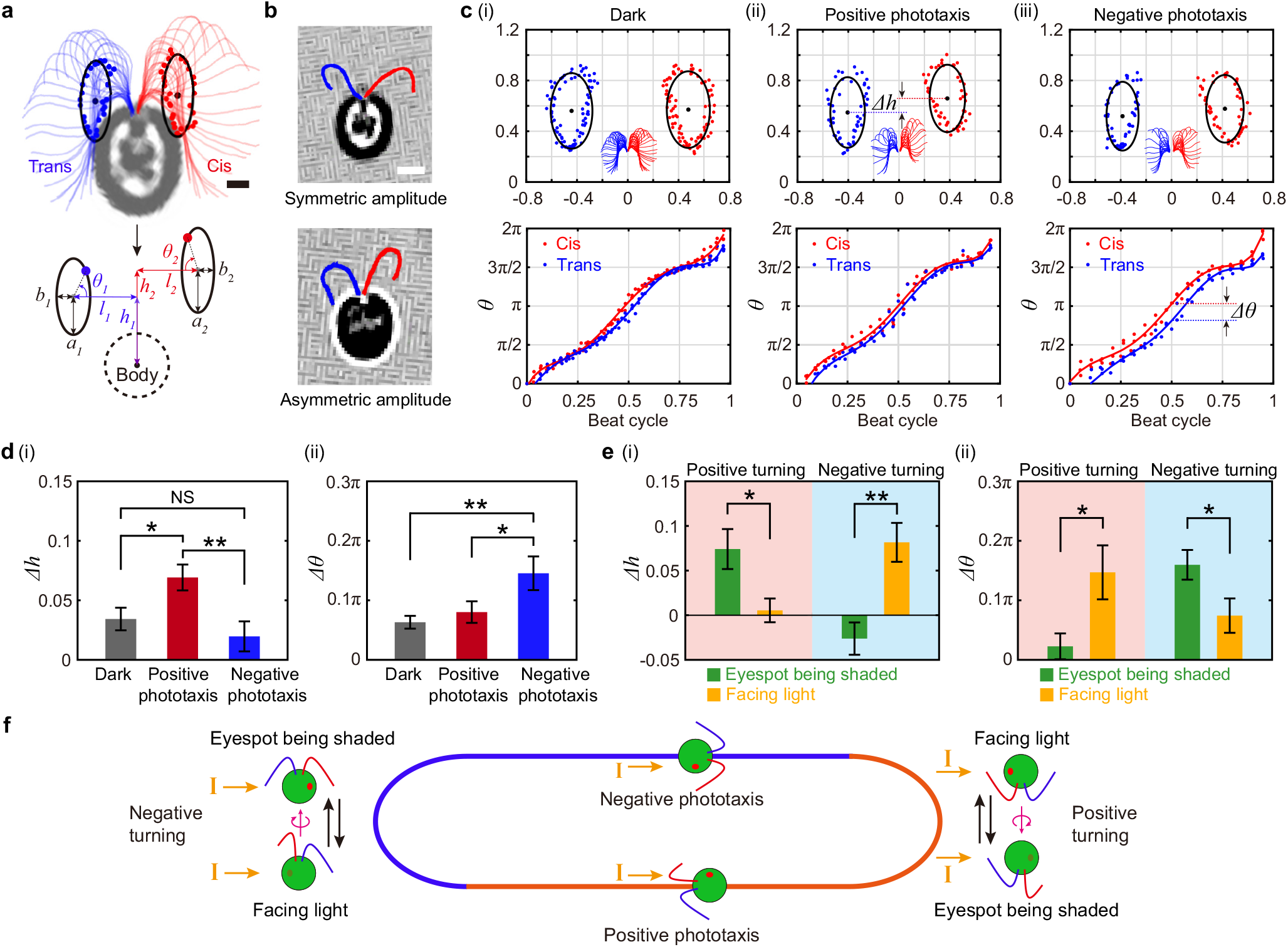
*Chlamydomonas* switch between symmetric and asymmetric flagella beats as well as modulating the flagella phase difference to transition between phototaxis modes. **a**, Flagella beat pattern was tracked manually and converted into elliptical orbits in the body reference frame, where the locations of flagella spheres correspond to the average locations of the two flagella outlines (Supplementary Section 2.6). **b**, Two major beat patterns with asymmetric beat amplitude and symmetric beat amplitude were observed in all phototaxis modes of *Chlamydomonas*. **c**, Flagella orbits (normalized by cell body length) and flagella phase over a beat cycle for cells under (i) darkness, (ii) positive phototaxis and (iii) negative phototaxis. The inset images represent the flagella waveforms for each case. The origin corresponds to the head of the cell where flagella attach to. **d**, (i, ii) Difference in beat amplitude Δ*h* and flagella phase difference Δ*θ* for cells at equilibrium states of positive and negative phototaxis as well as cells under darkness. Flagella beats were extracted when the beat plane was parallel to the light. Data were obtained from *n* = 6 cells with 3 beat cycles for each cell. **e**, Δ*h* and Δ*θ* for cells at turning states of positive and negative phototaxis. The green bars correspond to the case when the eyespot was shaded by cell body and no light is detected. The orange bars correspond to the case when the eyespot points towards the light. Data were obtained from *n* = 6 cells with 1 beat cycle for each cell. **f**, *Chlamydomonas* selects between symmetric and asymmetric beats according to light detection to switch between positive and negative phototaxis modes during oscillatory phototaxis. Scale bars, 3*µ*m (**a**) and 5*µ*m (**b**).

**Figure 4:**
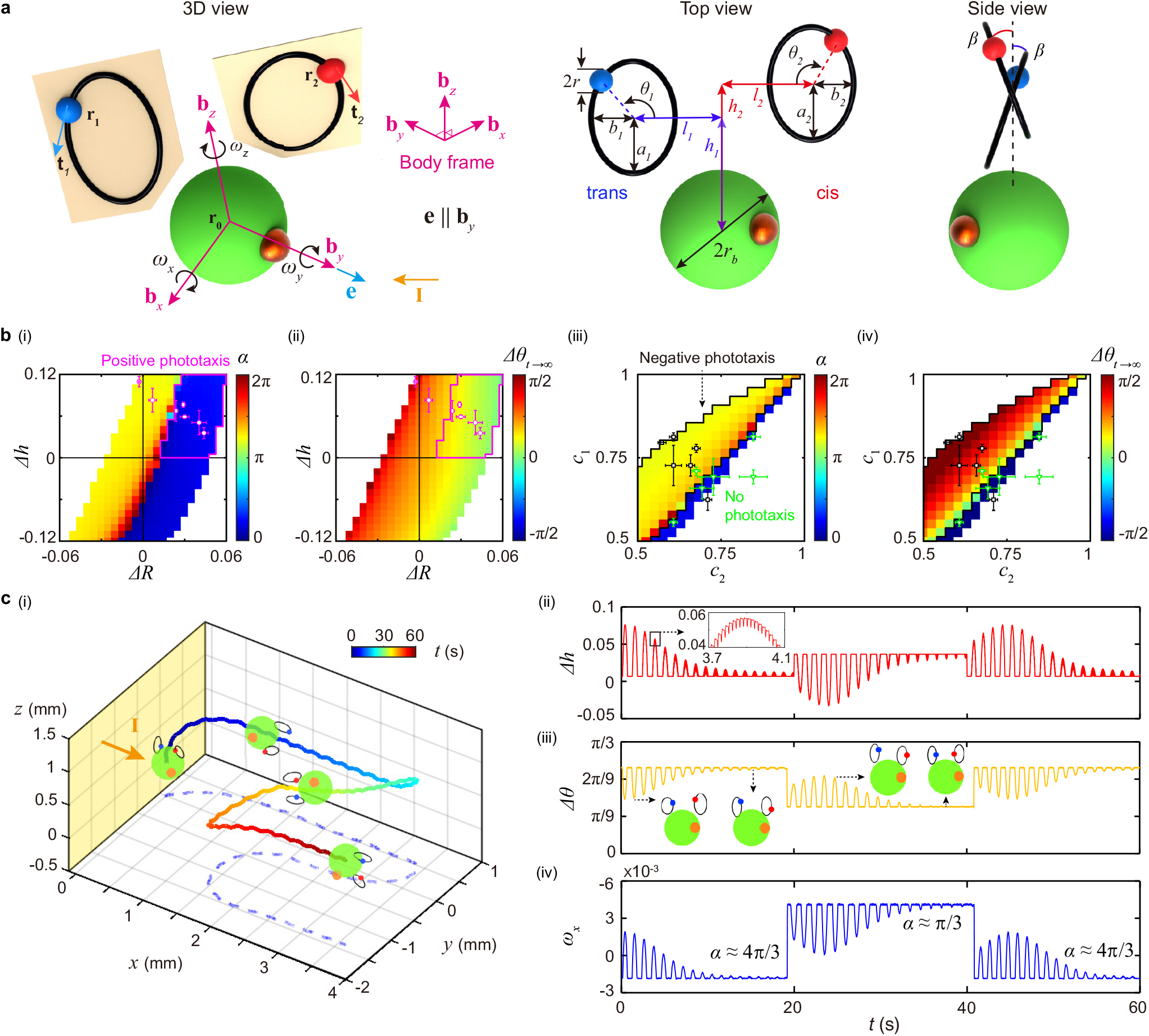
A photomechanical model captures oscillatory phototaxis of *Chlamydomonas*. **a**, Model description in 3D, top and side views. A pair of tilted beat orbits are considered in the model. **b**, (i,ii) Tuning of *α* and Δ*θ*_*t*→∞_ in the model by varying Δ*h* and Δ*R*, while keeping *c*_1_ and *c*_2_ constant. The region enclosed by purple lines represents the positive phototaxis regime in our simulations (0 *< α < π*). The purple markers denote Δ*h* and Δ*R* obtained from experimental data of positive phototaxis. (iii,iv) Tuning of *α* and Δ*θ*_*t*→∞_ by varying *c*_1_ and *c*_2_, where the same symmetric waveform was considered for all cases (Δ*h* = 0, Δ*R* = 0). The region enclosed by black lines represents the negative phototaxis regime in our simulations (*π < α <* 2*π*). The black markers and green markers denote *c*_1_ and *c*_2_ of simulations that fit the best with experimental phase angle and beat frequency of negative phototaxis and cells under darkness, respectively (Supplementary Section 4.3). **c**, Our model successfully captures the oscillatory phototaxis observed experimentally, showing quantitative agreement in the (i) trajectory and the modulation of (ii-iv) Δ*h*, Δ*θ* and *ω*_*x*_. The insets illustrate how the cell modulates its flagella orbits to modulate the parameters.

**Figure 5:**
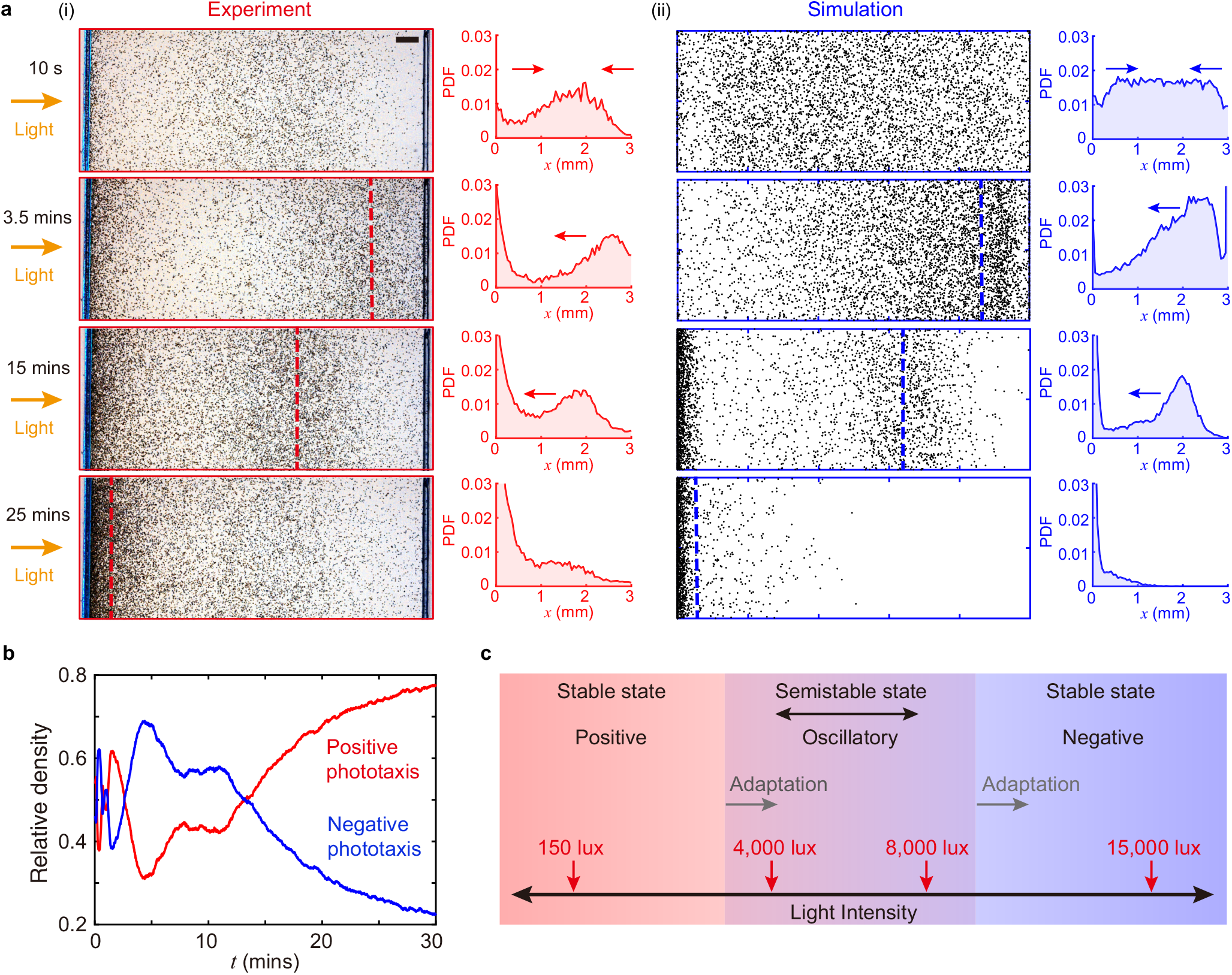
Adaptation of high density cells to light induces the emergence of a collective density wave. (**a**), Comparison of the propagation of density wave in experiment and simulation at several representative time instants. The probability density function (PDF) illustrates the cell distribution obtained by thresholding of images. The intensity of light stimulus is *∼* 8000 lux. Scale bar: 200*µ*m. We considered a cell number of 5000 in the simulation. **b**, Relative density of cells in the left half (red) and the right half (blue) of the chamber was measured as an approximation of the portion of cells exhibiting positive and negative phototaxis, respectively. The oscillation in relative density in the first *∼* 5 mins is due to cells alternating between positive and negative phototaxis. Afterwards, more and more cells adapt to light and transition to positive phototaxis completely, as can be seen from the increase (decrease) in relative density of positive (negative) phototaxis over time. **c**, A summary schematic elucidates the adaptation mechanism of *Chlamy-domonas*: The positive and negative phototaxis regime has an overlapping region which permits a semistable state of oscillatory phototaxis. The range of light intensities for oscillatory phototaxis shifts towards the negative phototaxis region over time, as indicated by the gray arrows, and more cells transition to positive phototaxis eventually.

### Phase angle tuning for oscillatory phototaxis

We first investigated how individual *Chlamydomonas* cells coordinate eyespot location and body orientation in different phototaxis modes (Fig. 2a). We tracked the cell orientation *φ* in the labora-tory frame (**x, y, z**) and the eyespot angle *ψ* in the body frame (**b**_*x*_, **b**_*y*_, **b**_*z*_). The helical swimming trajectory was described by 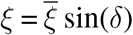. We obtained a fixed relationship between *δ* and *ψ* (Fig. 2b), in agreement to other microswimmers with helical swimming trajectories such as *E. gracilis* [5, 6]. This phase relationship was further captured by a phase angle *α* (Fig. 2b, Supplementary Section 2.4).

We experimentally measured *α* for positive phototaxis at weak light (*∼* 150 lux) and negative phototaxis at strong light (*∼* 15000 lux). We tracked *n* = 6 cells for each case and obtained *α* = 0.74 *±* 0.15 (mean*±*sem, throughout the paper if not stated otherwise) for positive phototaxis and *α* = 4.45 *±* 0.12 for negative phototaxis. We also observed *α* = 0.09 *±* 0.19 for cells under darkness (no phototaxis, *∼* 0 lux). These measured *α* for different phototaxis modes were in agreement with the prediction of our purely geometric phototaxis model (*P <* 0.0001, Fig. 2c, Supplementary Section 3), which showed 0 *< α < π* for positive phototaxis and *π < α <* 2*π* for negative phototaxis. This result agreed with previous reports which observed the eyespot pointing outward or inward of the helix for positive or negative phototaxis [18, 34]. Here, we further demonstrated that the eyespot, helical swimming and phototaxis modes are coupled with distinct phase relationships that can be quantified by a single parameter *α*.

We then tracked *α* for oscillatory phototaxis at intermediate light intensities (*∼* 4000 *−* 8000 lux). We divided the oscillatory trajectories into positive and negative portions when cells swam towards and away from light, respectively (Supplementary Section 2.5). We observed that *Chlamydomonas* changed *α* dynamically, where a relative swimming period was defined to normalize the time intervals for measuring *α* in a cycle (Fig. 2d). *α* for positive portions was consistent with positive phototaxis (*α* = 1.89 *±* 0.36), while *α* for negative portions appeared to split into two groups in the first and second halves of the relative swimming period (Fig. 2e). The first group had *α* = 4.54 *±* 0.30 (consistent with negative phototaxis) and the second group had *α* = 2.24 *±* 0.44 (consistent with positive phototaxis), indicating cells had an early transition from negative photo-taxis to positive phototaxis. This discrepancy can be explained as follow: Cells initially switched from negative to positive phototaxis had the eyespot pointed inward of the helix and away from light, avoiding them to make sharp turns immediately. In contrast, the eyespot pointed directly towards light when cells switched from positive to negative phototaxis and the cells made immediate turns. We also observed an increase in helix amplitude and beat frequency when cells transitioned from positive phototaxis to negative phototaxis (Supplementary Movie S4, Supplementary Section 2.8). The overall averaged *α* for oscillatory phototaxis was also significantly different from positive phototaxis (*P <* 0.05) and negative phototaxis (*P <* 0.001), indicating that it is indeed a distinct phototaxis mode (Fig. 2f). We conclude that oscillatory phototaxis emerges from the continuous switching between positive and negative phototaxis via dynamic tuning of *α*.

### Flagella beat patterns for switching of phototaxis modes

To reveal how *Chlamydomonas* coordinate flagella beats to tune their *α*, we quantitatively tracked the flagella at equilibrium state after completing turns (Fig. 1b (ii & iv)) as well as the flagella at turning state (Fig. 1b (iii & v)). We manually tracked the flagellum outlines (*n* = 6 cells for 3 beat cycles for each phototaxis mode, *∼* 20 frames per cycle, sampling rate 1000 fps) and converted the beat patterns into elliptical orbits (Fig. 3a, Supplementary Section 2.6). *a*_*i*_ and *b*_*i*_ denote the orbit axes. *h*_*i*_ and *l*_*i*_ denote the distances between orbit centers and body center (*i* = 1, 2 for trans- and cis-flagellum). The flagella phases are defined as *θ*_*i*_. It turns out that the switching between phototaxis modes can be captured by selection of two distinct flagella beat patterns (Fig. 3b, Supplementary Movie S5) characterized by symmetric (Δ*h* = *h*_2_ *− h*_1_ *≈* 0) and asymmetric amplitudes (Δ*h >* 0), while the change in Δ*l* = *l*_2_ *− l*_1_ is not obvious.

We first compared the flagella beats for cells under darkness and equilibrium state of positive and negative phototaxis (Fig. 3c). We observed symmetric beats for cells under darkness and asymmetric beats for positive phototaxis (Fig. 3c (i) & (ii)), with a significant difference in Δ*h* (Fig. 3d (i), *P <* 0.01, *n* = 6). Yet, there was no significant difference in Δ*h* for negative phototaxis and cells under darkness (Fig. 3c (i) & (iii), Fig. 3d (i)), but instead we observed a significant larger flagella phase difference (Δ*θ* = *θ*_2_ *− θ*_1_) for negative phototaxis (Fig. 3d(ii), *P <* 0.01, *n* = 6).

We then compared the flagella beats at turning state and we again observed two beat patterns similar to equilibrium state when the eyespot faces light or being shaded in a roll cycle (Fig. 3e; Supplementary Movie S6, S7). For positive phototaxis, the cells switched from asymmetric beats when eyespot being shaded to symmetric beats when eyespot faced light, where Δ*h* decreased (Fig. 3e(i)) and Δ*θ* increased (Fig. 3e(ii)). The opposite switching in beat patterns was observed for negative phototaxis. Thus, the beat patterns observed when eyespot being shaded at turning state were consistent with beat patterns at equilibrium state, which in both situations the eyespot did not detect light effectively.

These results lead to a comprehensive understanding of the phototaxis mechanism (Fig. 3f): *Chlamydomonas* can dynamically tune the phase angle *α* to achieve positive and negative phototaxis by selecting between symmetric and asymmetric beats (by tuning Δ*h*) as well as modulating the phase difference (Δ*θ*) between the cis- and trans-flagellum. The cell shifts to the other beat pattern when its eyespot detects a light stimulus. The beat pattern is recovered when the eyespot is being shaded or when the cell reaches the equilibrium state.

### Hydrodynamic model for oscillatory phototaxis

To further verify the proposed phototaxis mechanism, we implement a coarse-grained hydrodynamic model extended from three-sphere models proposed previously (Fig. 4a, Supplementary Section 4) [17, 35, 36]. The model consists of a large cell body sphere with radius *r*_*b*_ located at **r**_0_ and two smaller flagella spheres with radius *r* located at **r**_*i*_ (*i* = 1, 2 for trans- and cis-flagellum, *r* ≪ *r*_*b*_). The flagella spheres are constrained to move on elliptical orbits tilted by an angle *β* (Fig. 4a), which was measured to be 0.3 rad [17]. However, this reduced model would underestimate the effective forces generated by the flagella [17]. Thus, we scaled the orbit parameters to account for the mismatch (Supplementary Section 4.1.3).

The flagella spheres are driven by tangential forces 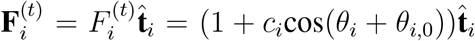 along the orbits, where *c*_*i*_ and *θ*_*i*,0_ are constants accounting for the amplitudes of force modulation and the initial force phases of the cis- and trans-flagellum. The sphere velocities can be determined by their hydrodynamic interactions via the Oseen approximation [37]:

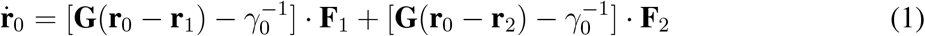

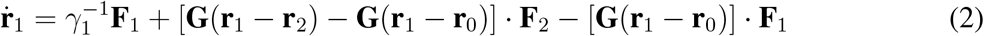

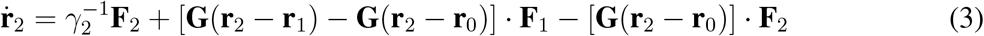

where *γ*_*i*_ = 6*πηr, γ*_0_ = 6*πηr*_*b*_ and 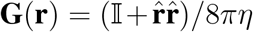. Here *η* is the fluid dynamic viscosity. Together with the force-free and torque-free conditions, the motions of cell body and flagella spheres can be fully solved, including the angular velocity of the cell body (*ω*_*x*_, *ω*_*y*_, *ω*_*z*_). The phase angle can be calculated as *α* = tan^*−*1^(*ω*_*y*_*/ω*_*x*_).

The photoresponse of *Chlamydomonas* is modeled by linking the light hitting the eyespot to the flagella beat pattern. The light signal *S*(*t*) detected by the cell is given by

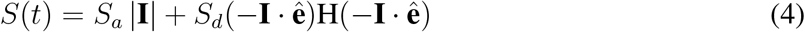

Here **I** and 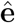 are the light vector and the eyespot vector (Fig. 4a). *S*_*a*_ and *S*_*d*_ are the coupling constants for ambient light and directional light. H is the Heaviside function accounting for shading of light. The cell modulates the flagella beat pattern according to the detected light: *h*_*i*_ = *h*_0_ + *K*_*h,i*_ log(*S*(*t*)) and *R*_*i*_ = *R*_0_ + *K*_*R,i*_ log(*S*(*t*)), where *R*_*i*_ = (*a*_*i*_ + *b*_*i*_)*/*2 is the beat amplitude and Δ*R* = *R*_2_ *− R*_1_. *K*_*h,i*_ and *K*_*R,i*_ are light-dependent coupling constants.

We verified the phototaxis mechanism by comparing the parameters for tuning of *α* from our model predictions and experimental data. We first varied Δ*h* and Δ*R* but keeping *c*_1_ = *c*_2_ in our model, which effectively tuned *α* by changing flagella waveform. We obtained phase maps for *α* and the flagella phase difference Δ*θ*_*t*→∞_ at the equilibrium state (Fig. 4b (i) and (ii), Supplementary Section 4.3). Most experimental data for positive phototaxis (purple markers) falls within positive phototaxis regime of the model (enclosed by purple border). A few outliers occur when Δ*R ∼* 0, corresponding to cases which the cis- and trans-flagellum have similar beat amplitudes, and the reduced model may be less accurate to capture the delicate difference in flagella forces. We then varied *c*_1_, *c*_2_ but keeping Δ*h ∼* 0 and Δ*R ∼* 0, which changed *α* by varying Δ*θ*_*t*→∞_ (Fig. 4b (iii) and (iv)). Nearly all experimental data for negative phototaxis (black markers) falls within the negative phototaxis regime (enclosed by black border). We also observed that the experimental data for cells under darkness (green markers) correspond to *α ∼* 0 in our model which no phototaxis occurs. Further discussions on other parameters such as beat frequency and helix diameter can be found in Supplementary Section 4.3. We further simulated the oscillatory phototaxis (Fig. 4c(i), Supplementary Movie S8-S10) and examined the modulation of Δ*h*, Δ*θ*, and *ω*_*x*_ (Fig. 4c(ii))-(iv)). In the negative portions of the oscillatory phototaxis, we observed Δ*h ∼* 0 and larger Δ*θ* at the equilibrium state and Δ*h >* 0 and smaller Δ*θ* at the turning state, agreeing with experimental observations (Fig. 3d). The same conclusion can be made for positive portions. Hence, our model captures quantitatively the transition between different phototaxis modes (Supplementary Movie S11, S12).

Several other hypotheses of phototaxis mechanisms for *Chlamydomonas* were proposed based on the dominance of trans- or cis-flagellum or modulation of flagella tangential force [15–18]. Our model can capture all these other possible phototaxis mechanisms. A full discussion of the modulation of different orbit parameters can be found in Supplementary Section 4.4. Yet, we verify experimentally that switching of flagella waveforms and flagella phase modulation are the key mechanisms for phototaxis transition. Our unified phototaxis mechanism resolves the long-standing argument of how *Chlamydomonas* transition between positive and negative phototaxis.

### Adaptation mechanism for density wave formation

Finally, we verified how adaptation over multiple time scales induces the emergence of density wave at the population level (Fig. 5, Supplementary Section 5). The propagation of density wave was captured via thresholding of cell distribution (Fig. 5a). At short times (*∼* 1 *−* 2 s), adaptive flagella responses to light tuned *α* instantaneously to determine phototaxis sign (Fig. 4c). After *∼* 10 s of light exposure, density distribution rapidly oscillated as cells split into two groups with positive and negative phototaxis initially (data obtained from individual tracking of *n* = 32 cells, Supplementary Section 5.1). At longer times (*∼*1 min), cells switched phototaxis sign based on intracellular biochemical signals, influenced by factors such as intracellular redox state [12] and phosphorylation of channelrhodopsin [38]. Two rhodopsins in *Chlamydomonas reinhardtii* were found to mediate phototaxis at low and high light intensities with different adaptation rates, respectively [23, 39]. This adaptation rate discrepancy may change the dominant rhodopsin and switch phototaxis sign. We observed a distribution of swimming periods from different cells during oscillatory phototaxis (Supplementary Section 5.1), indicating a diversity in adaptation rates. Density wave emerged at *∼* 3.5 mins from the peak distribution of cells having similar adaptation rates. At even longer times (*>* 15 mins), more cells transitioned to positive phototaxis due to long-term adaptation (Fig. 5b). The wave gradually traveled towards the light source and reached the near-light side of the chamber at *∼* 25 *−* 30 mins when most cells completely adapted to light and exhibited only positive phototaxis.

To further explain the formation of density wave, we develop a general adaptation model that captures accumulation and relaxation of biochemical signals for determining phototaxis sign (model details can be found in Supplementary Section 5):

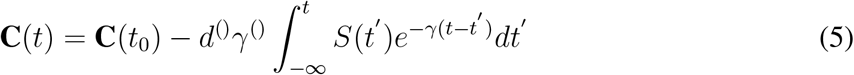

Here **C**(*t*) varies between 0 and 1 to switch between positive phototaxis (*d*^+^ *<* 0) and negative phototaxis (*d*^*−*^ *>* 0), respectively. The time-dependent adaptation rates (*γ*^+^ and *γ*^*−*^) are obtained by fitting swimming periods of tracked cells during oscillatory phototaxis (Supplementary Section 5.2).

The adaptation model was sufficient to capture the emergence of density wave, but we observed a mismatch in length and time scales of the simulated density wave with experiments. We therefore incorporated additional effects including speed reduction due to cell collision at high density (cell-cell interactions not directly considered) and cell adhesion/detachment on glass surface as reported previously [26] (Supplementary Section 5.3, Supplementary Movie S13). The resulting simulated density wave closely resembled experimental observation (Fig. 5b, Supplementary Movie S14).

## Discussion

We conclude that oscillatory phototaxis is a semi-stable state due to overlapping light intensity thresholds for positive and negative phototaxis (Fig. 5c). These phototaxis thresholds shift over times due to adaptation and cells transition to positive phototaxis eventually. Oscillatory phototaxis provides a mechanism for *Chlamydomonas* to avoid moving too far away from light by turning back occasionally and avoid photodamage if the light is still too strong after turning, effectively expanding the survival range of light intensity for processes such as photosynthesis and cell division. Moreover, individual cells within a density wave can act as effective light shelters and diffusers to provide collective photoprotection for the colonies. A similar photoprotection mechanism was observed in light-induced phase separation of *Chlamydomonas* [40], where cells formed branching patterns instead of a density wave. The mechanisms deciphered for phototaxis transition and density wave formation in this work promise generalizable biophysical laws for microswimmer navigation [6, 41–44] and inform the control of collective density waves emerged in active matter systems [45–49].

## Acknowledgement

This work was supported by the Research Grants Council of Hong Kong through the General Research Fund (No. 27208421 and No. 17303423) and the Croucher Foundation. We thank members of the Tsang lab and Riedel-Kruse for discussions.

